# LRR-extensins of vegetative tissues are a functionally conserved family of RALF1 receptors interacting with the receptor kinase FERONIA

**DOI:** 10.1101/783266

**Authors:** Aline Herger, Shibu Gupta, Gabor Kadler, Christina Maria Franck, Aurélien Boisson-Dernier, Christoph Ringli

## Abstract

Plant cell growth requires the coordinated expansion of the protoplast and the cell wall that confers mechanical stability to the cell. An elaborate system of cell wall integrity sensors monitors cell wall structures and conveys information on cell wall composition and growth factors to the cell. LRR-extensins (LRXs) are cell wall-attached extracellular regulators of cell wall formation and high-affinity binding sites for RALF (rapid alkalinization factor) peptide hormones that trigger diverse physiological processes related to cell growth. RALF peptides are also perceived by receptors at the plasma membrane and LRX4 of *Arabidopsis thaliana* has been shown to also interact with one of these receptors, FERONIA (FER). Here, we demonstrate that several LRXs, including the main LRX protein of root hairs, LRX1, interact with FER and RALF1 to coordinate growth processes. Membrane association of LRXs correlate with binding to FER, indicating that LRXs represent a physical link between intra- and extracellular compartments via interaction with membrane-localized proteins. Finally, despite evolutionary diversification of the LRR domains of various LRX proteins, many of them are functionally still overlapping, indicative of LRX proteins being central players in regulatory processes that are conserved in very different cell types.

**Author Summary:** Cell growth in plants requires the coordinated enlargement of the cell and the surrounding cell wall, which is ascertained by an elaborate system of cell wall integrity sensors, proteins involved in the exchange of information between the cell and the cell wall. In *Arabidopsis thaliana*, LRR-extensins (LRXs) are localized in the cell wall and are binding RALF peptides, hormones that regulate cell growth-related processes. LRX4 also binds the plasma membrane-localized receptor kinase FERONIA (FER), establishing a link between the cell and the cell wall. It is not clear, however, whether the different LRXs of Arabidopsis have similar functions and how they interact with their binding partners. Here, we demonstrate that interaction with FER and RALFs requires the LRR domain of LRXs and several but not all LRXs can bind these proteins. This explains the observation that mutations in several of the *LRXs* induce phenotypes comparable to a *fer* mutant, establishing that LRX-FER interaction is important for proper cell growth. Some LRXs, however, appear to influence cell growth processes in different ways, which remain to be identified.

## Introduction

The plant cell wall is a complex structure of interwoven polysaccharides and structural proteins that protects the cell from biotic and abiotic stresses [1]. Importantly, it serves as a shape-determining structure that resists the internal turgor pressure emanating from the vacuole. Cell growth requires a tightly regulated expansion of the cell wall, generally accompanied by the concomitant biosynthesis of new cell wall material that is integrated into the expanding cell wall [2]. The signal transduction machinery required for coordinating the intra- and extracellular processes involves a number of transmembrane proteins at the plasma membrane to connect the different cellular compartments [3]. Among those, the *Catharanthus roseus* receptor-like kinase 1-like protein (*Cr*RLK1L) THESEUS1 revealed to be a cell wall integrity sensor that perceives reduced cellulose content in the cell wall and induces compensatory changes in cell wall composition to restrain growth [4]. Several members of the *Cr*RLK1L family are involved in cell growth processes [5,6,7,8,9]. FERONIA (FER) is required for proper pollen tube reception during the fertilization process involving local disintegration of the cell wall [10]. The extracellular domain of FER has been demonstrated to bind pectin, a major component of cell walls [11]. This interaction could contribute to the function of FER during cell wall integrity sensing and perception of mechanical stresses [11,12,13]. Several *Cr*RLK1L receptors have been demonstrated to bind rapid alkalinization factor (RALF) peptides, that induce alkalinization of the extracellular matrix, change Ca2+ fluxes and modulate cell growth and response to pathogens [14,15,16,17,18,19,20,21]. Hence, CrRLK1L proteins appear to have multiple functions, suggesting that their activity is at the nexus of different cell growth-related activities.

RALF1 was identified as a ligand of FER and a number of proteins are involved in the RALF1-FER triggered signaling process, either as signaling intermediates such as ROP2, ROPGEF, ABI2, RIPK [22,23,24], co-receptor such as BAK1 [25], or as targets of the FER-dependent pathway, such as AHA2 [15]. The receptor kinase-like protein MARIS (MRI) and the phosphatase ATUNIS1 (AUN1) were identified as downstream components of signaling activities induced by ANXUR1 and 2 (ANX1/2), pollen-expressed FER homologs [6,26,27]. It is not clear at this point, however, to what extent the signaling components are shared among the different CrRLK1Ls. Both AUN1 and MRI also influence root hair growth, indicating that they might function downstream of a root hair-expressed *Cr*RLK1L protein [26].

LRX (LRR-extensins) are extracellular proteins involved in cell wall formation and cell growth. They consist of an N-terminal (NT) domain and a Leucine-rich repeat (LRR) of 11 repeats, followed by a short Cys-rich domain (CRD) serving as a linker to the C-terminal extensin domain (Figure 1) [28,29]. The extensin domain contains Ser-Hyp_n_ repetitive sequences that are characteristic for hydroxyproline-rich glycoproteins[30,31] and appears to serve in anchoring the protein in the extracellular matrix [32,33]. The N-terminal moiety with the NT- and LRR-domain associates with the membrane fraction [34], indicating a function of LRXs in linking the cell wall with the plasma membrane by binding of a membrane localized interaction partner. The recent identification of LRX4 as an interactor of FER corroborates this hypothesis [35].

**Figure 1.**
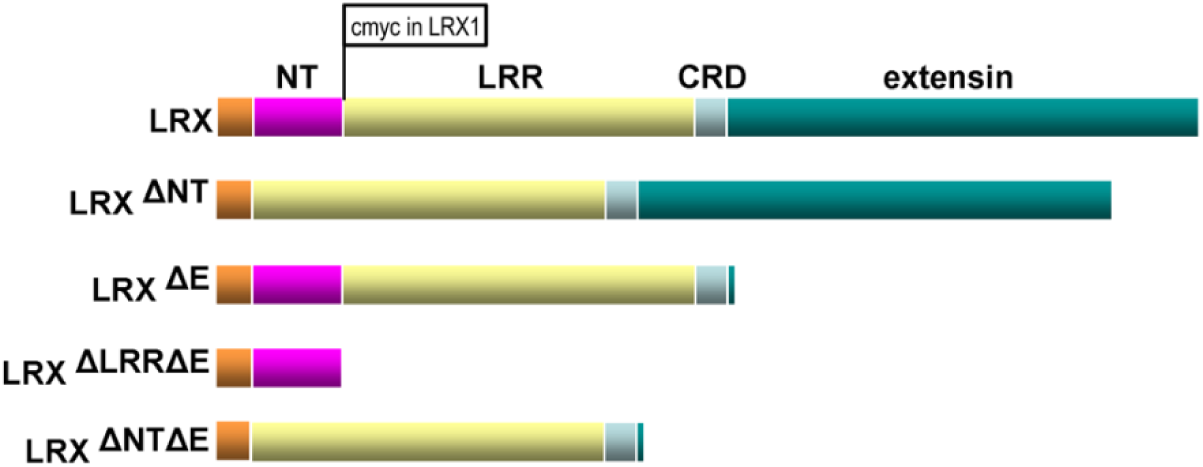
Structure of LRX proteins and deletion constructs used in this study. LRX proteins consist of a signal peptide for export of the protein (light brown), an NT-terminal domain (purple) of unknown function, an leucine-rich repeat (LRR) domain (yellow), a Cys-rich hinge region (CRD), and a C-terminal extensin domain (green), with Ser-Hyp_n_ repeats typical of hydroxyproline-rich glycoproteins, for insolubilization of the protein in the cell wall. The different deletion constructs used in this study are listed, with “Δ” indicating deleted domains. In the LRX1 construct, a cmyc tag was introduced between the NT- and the LRR-domain, which does not interfere with protein function and allows for immuno-detection of LRX1.

The *LRX* family of Arabidopsis consist of eleven members, most of which are expressed in a tissue-specific manner. *LRX1/2, LRX3/4/5,* and *LRX8/9/10/11* are predominantly expressed in root hairs, in the main root and the shoot, and pollen, respectively. Mutations in these genes cause cell wall perturbation and cell growth defects in the respective cell types [32,34,36,37]. *lrx1* mutants develop deformed root hairs that are swollen, branched, and frequently burst [32]. This phenotype is strongly enhanced in *lrx1 lrx2* double mutants that are virtually root hair less [36]. The *lrx345* triple mutant shows defects in vacuole development, monitoring of cell wall modifications, and sensitivity to salt stress that are reminiscent of a *fer* mutant [35,38], which is in line with LRX4 and possibly other LRXs interacting with FER. Mutants affected in several of the pollen-expressed *LRXs* are impaired in pollen tube growth and show reduced fertility [34,39,40].

LRX proteins were recently identified as high-affinity binding sites for RALF peptides, with the binding spectrum differing among the LRXs. LRX8 of pollen tubes was shown to physically interact with RALF4 [41] while in vegetative tissues, the root/shoot-expressed LRX3, LRX4, and LRX5 were reported to bind RALF1,22,23,24, and 31 [38,42]. Whether and how the binding of RALFs to LRXs and FER influence the interaction of these proteins remains to be investigated. It is also not clear to what extent the different LRXs are functionally similar and whether they share FER as a common interaction partner.

Here, we analyzed LRX protein functions and demonstrate that the membrane-association of LRXs correlates with the ability to bind FER. The root hair-expressed LRX1 binds FER and RALF1, and this binding activity is also revealed for other LRX proteins. Together with the *fer*- like phenotype of higher-order *lrx* mutants, this suggests that LRX proteins of different vegetative tissues interact with the ubiquitously expressed FER. Cross-complementation experiments of *lrx* mutants suggest that some but not all LRX proteins exert similar functions. Together with recently published data, this suggests that LRX proteins interact with RALF peptides and FER, but that they also carry additional functions independent of these protein-protein interactions that are relevant for the regulation of cell growth.

## Results

### The membrane association and interaction of LRX4 with FER dependents on the LRR domain

We have previously shown that LRX proteins associate with the membrane and LRX4 binds the *Cr*RLK1L FER [34,42]. LRX4 deletion constructs were produced to test for correlation between FER binding and membrane association. The *LRX4* promoter was used to express *LRX4*^ΔE^*-HA* (coding for LRX4 missing the extensin domain), *LRX4^ΔLRRΔE^-HA* (coding for LRX4 missing the LRR- and the extensin domain), and *LRX4 ^ΔNTΔE^-HA* (coding for LRX4 missing the NT- and the extensin domain) (Figure 1) in transgenic Arabidopsis. (Gene identifiers of all genes used in this study are listed in the Material and Methods section.) Extensin deletion constructs were used in this and in later experiments to prevent insolubilization of the LRX protein in the cell wall. Membrane fractions of the different transgenic lines were isolated from seedlings and tested for presence of the recombinant protein. As shown in Figure 2A, all proteins were present in the total fraction. While LRX4^ΔE^-HA and LRX4^ΔNTΔE^-HA were also detected in the membrane fraction, LRX4^ΔLRRΔE^-HA was not. Successful isolation of membrane fractions was confirmed by detection of the membrane-marker protein LHC1a [43]. This demonstrates that the membrane association of LRX4 depends on the presence of its LRR domain. Next, the constructs *LRX4^ΔNTΔE^-HA* and *LRX4^ΔLRRΔE^-HA* were expressed under the *35S CaMV* promoter (subsequently referred to as *35S*) in *N. benthamiana* for co-immunoprecipitation (Co-IP) experiments with the extracellular domain (ECD) of FER fused to citrine (FER^ECD^-citrine) or FLAG (FER^ECD^-FLAG). Co-IP analysis revealed that the FER^ECD^ was co-purified when expressed with LRX4^ΔNTΔE^-HA (Figure 2B) but not with LRX4^ΔLRRΔE^-HA (Figure 2C). These analyses reveal a positive correlation between LRX4*^ΔE^* binding to FER and its association with the plasma membrane.

**Figure 2.**
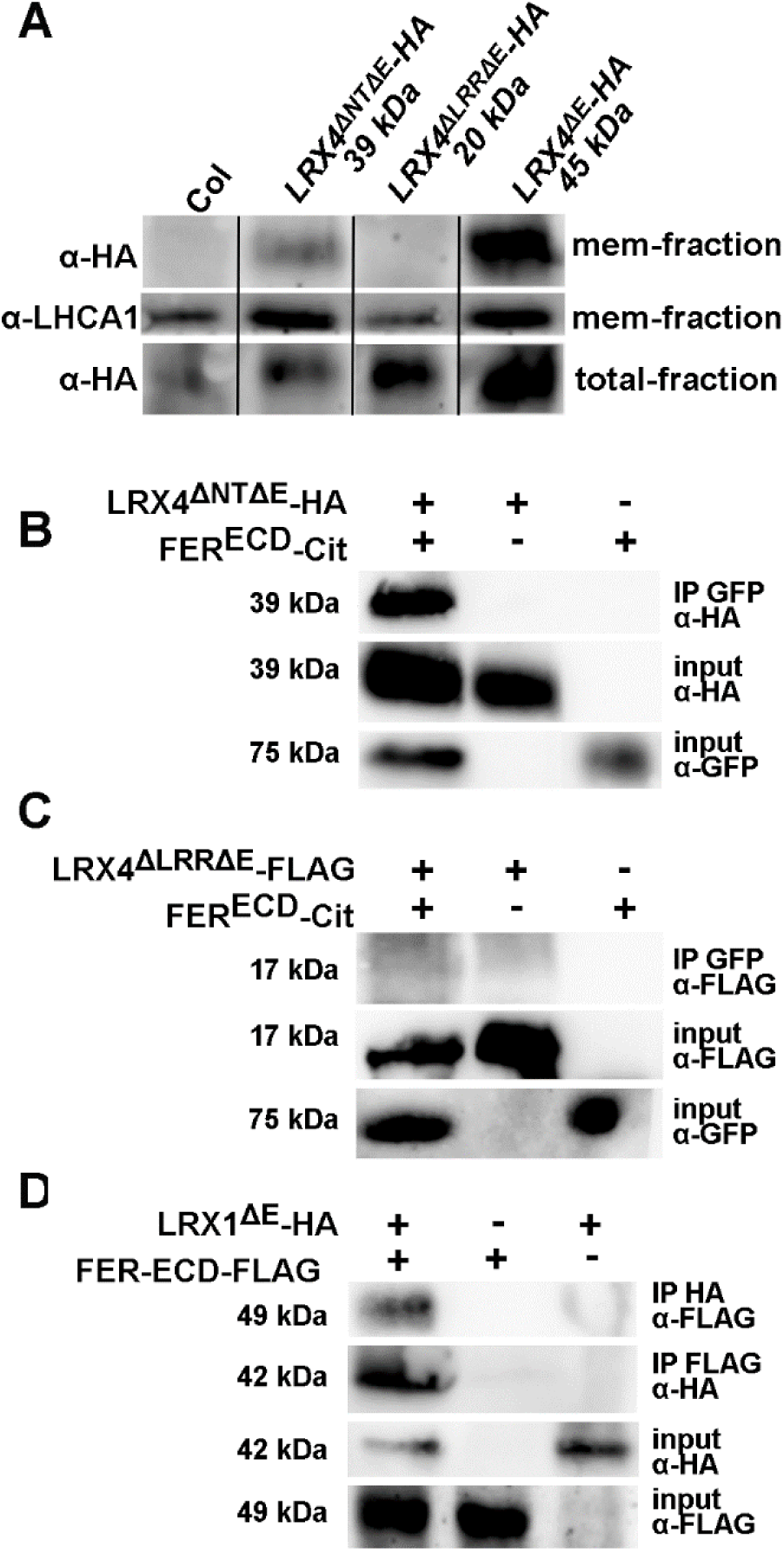
Correlation of membrane association and interaction with FER. (A) Western-blot of membrane fractions of transgenic Arabidopsis expressing constructs as indicated. LHCA1 is a membrane-associated protein confirming identity of membrane fraction. (B and C) LRX4^ΔNTΔE^ (B) but not LRX4^ΔLRRΔE^ (C) can be co-purified with FER^ECD^ when expressed in tobacco, indicating that the LRR domain is sufficient and necessary for in planta interaction with FER^ECD^. (D) When co-expressed in tobacco LRX1^ΔE^ and FER^ECD^ can be co-immunoprecipitated, indicative for the interaction of the two proteins. Antibodies used for IP and subsequent detection by western blotting are indicated.

### Several LRXs of vegetative tissue interact with FER

The root hair-expressed LRX1 is so far the best characterized LRX protein and the *lrx1* root hair mutant represents a convenient genetic system for analyses of LRX protein function [32,33,44,45]. Since FER was reported to maintain cell wall integrity in growing root hairs [23,26], it was interesting to test whether LRX1 also interacts with FER. To this end, constructs encoding LRX1^ΔE^-HA and FER^ECD^-citrine under the *35S* promoter were expressed in tobacco for Co-IP experiments. LRX1^ΔE^ shows interaction with FER^ECD^ (Figure 2D). Interaction of LRX1 with FER^ECD^ was also confirmed in a yeast-two-hybrid experiment (Suppl. Figure S1). The yeast-two-hybrid experiments were extended to other LRXs of vegetative tissues, namely LRX2, LRX3, LRX4, and LRX5. While LRX2, LRX4, and LRX5 showed interaction with FER^ECD^, LRX3 failed to interact (Suppl. Figure S1). However, the BD-LRX3 did not accumulate to detectable levels in yeast extracts (data not shown). Hence, a conclusion on LRX3-FER^ECD^ interaction cannot be drawn. Therefore, Co-IP experiments were conducted with LRX3^ΔE^ and FER^ECD^, but failed to show interaction of these two proteins, as found by others [38].

### The *lrx12345* quintuple mutant mimics the *fer-4* mutant phenotype

The results obtained above suggest that the five LRX proteins expressed in vegetative tissue that have been analyzed so far could exert overlapping functions. Since a double mutant for the root hair-expressed *LRX1* and *LRX2* [36] displays a root hair phenotype comparable to the knock-out mutant *fer-4* (Duan et al., 2008), and the *lrx345* triple mutant develops a shoot phenotype that is reminiscent of *fer-4* [38,42], we anticipated that an *lrx12345* quintuple mutant would be globally similar to *fer-4*. The *lrx1 lrx2* mutant was crossed with the *lrx345* triple mutant and an *lrx12345* quintuple mutant was identified in the segregating F2 population of this cross based on a root hair-less root and retarded shoot growth with an increase in anthocyanin content (Figure 3A). Indeed, the *lrx12345* quintuple mutant shows *fer-4* like phenotypes in the root and shoot at the seedling stage and, at the adult stage, smaller and broader rosette leaves with increased accumulation of anthocyanin compared to the wild type (Figure 3A). *fer-4* seedlings grown in vertical orientation display reduced gravitropic growth of the root [46]. This growth defect was assessed in the wild type, *fer-4*, and different *lrx* mutant combinations by assessing the vertical growth index [47]. For quantification, the ratio between the absolute root length and the progression of the root along the gravity vector, the arccos of α, was used as illustrated in Figure 3B. Accumulating *lrx* mutations cause an agravitropic response comparable to *fer-4* (Figure 3C). Thus, the genetic analysis of higher-order *lrx* mutants and *fer-4* support the finding that most of the LRXs are active in the signaling pathway of FER and are able to interact with FER.

**Figure 3.**
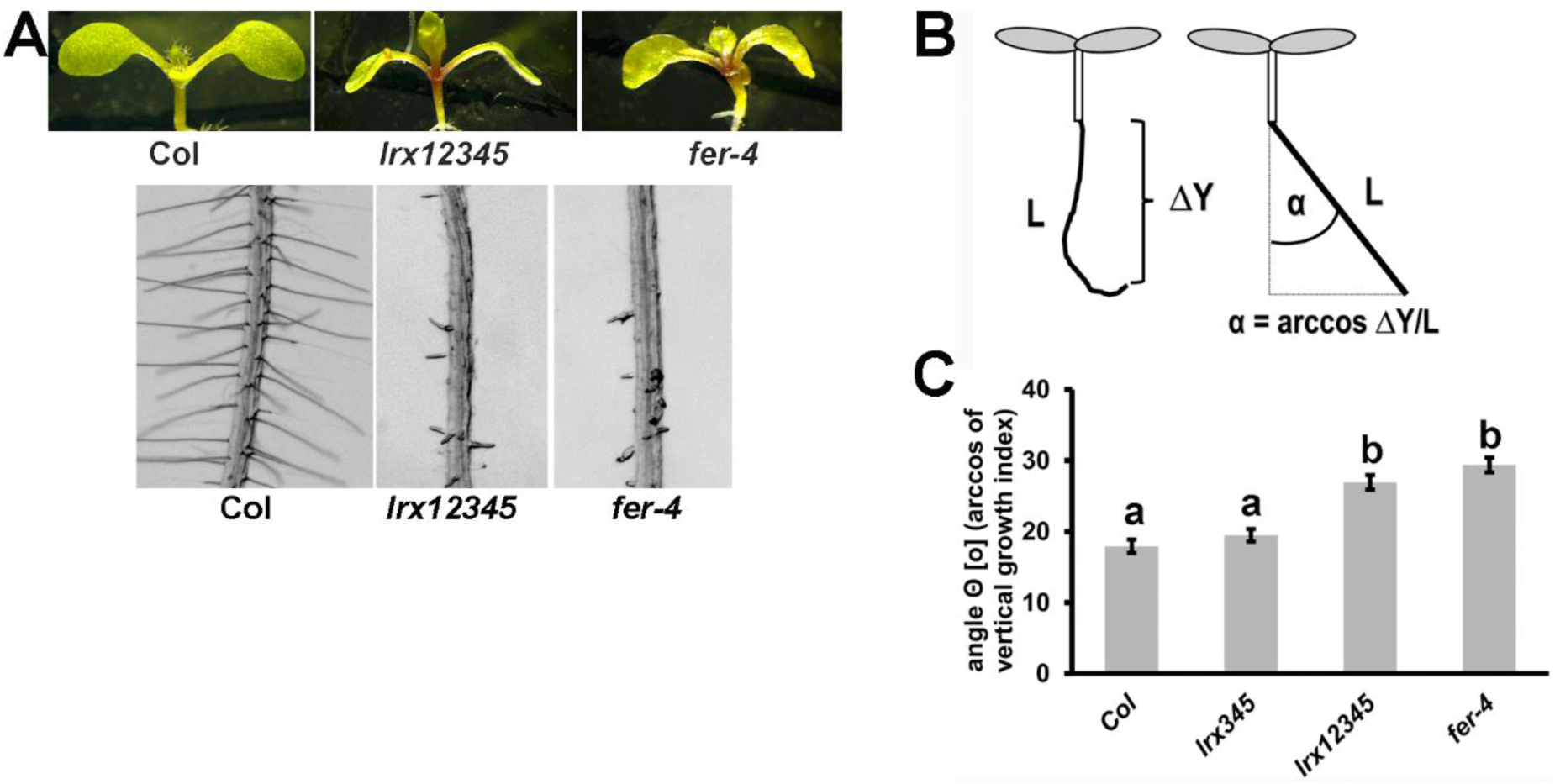
*fer-4* and *lrx12345* mutants show comparable phenotypes. (A) Seedlings were grown for 5 days on half-strength MS for analysis of root hair formation (bottom) and another 5 days for analysis of shoot development (top). (B) Quantification of gravitropic response by the root growth index. (C) Increasing agravitropy by accumulation of *lrx* mutations. Error bars represent SEM. Different letters above bars indicate significant differences (T-test, n>20, P<0.0001). Bar=5mm (A, top lane), 0.5 mm (A bottom lane)

### The LRR domain is necessary for the dominant negative effect of LRX1 missing the extensin domain

Expression of a truncated version of LRX1 lacking the extensin coding sequence (LRX1^ΔE^; Figure 1) under the *LRX1* promoter induces a dominant-negative effect in wild-type seedlings, resulting in a defect in root hair formation [32,33], possibly because LRX1^ΔE^ competes with the endogenous LRX1 for binding partners. This observed activity of LRX1^ΔE^ was used for further functional analysis of the LRX1 protein. Specifically, we assessed which domains are required or dispensable for the dominant negative effect. Of the LRX1^ΔE^ construct, the LRR domain or the NT domain were removed, resulting in LRX1^ΔLRRΔE^ and LRX1^ΔNTΔE^, respectively (Figure 1). The corresponding constructs under the *LRX1* promoter were transformed into wild-type Columbia plants. T2 seedlings expressing either of the two constructs developed wild-type root hairs (Figure 4A), hence failed to produce the dominant-negative effect on root hair development. Extracts from root tissue of the different lines were used for western blotting. An antibody detecting the cmyc-tag of the recombinant LRX1 variants confirmed that the proteins were produced. As shown in Figure 4B, the transgenic lines produce proteins with the expected decrease in mass of LRX1^ΔE^ > LRX1*^ΔNT^*^ΔE^ > LRX1*^ΔLRR^*^ΔE^. Together, this indicates that both the LRR- and the NT-domain are required but neither is sufficient to induce the dominant-negative effect on root hair development.

**Figure 4.**
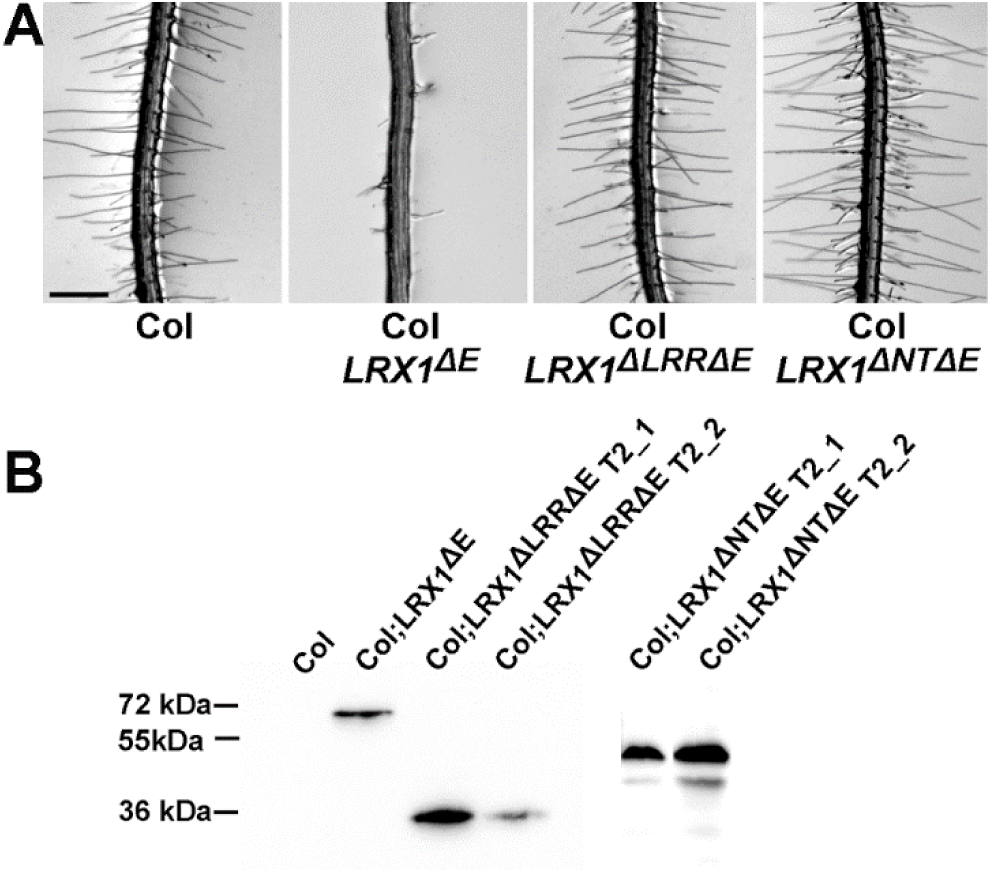
Both the LRR and NT domains are required for inducing the dominant negative effect of LRX1^ΔE^. (A) Wild type seedlings (Col) or transgenic Col lines expressing LRX1^ΔE^, LRX1^ΔLRRΔE^, or LRX1^ΔNTΔE^ (for structure see Figure 1). The dominant negative effect of LRX1^ΔE^ depends on both the LRR and the NT domains. A representative example of several independent transgenic lines is shown. (B) Western-blot using an anti-cmyc antibody detecting the proteins encoded by the transgenic lines shown in (A). Bar= 0.5 mm

In a complementary approach, we tested whether the NT-domain is required for the function of the full-length LRX1. To this end, the *lrx1* and *lrx1 lrx2* mutants developing intermediate and strong root hair defects, respectively [36], were transformed with the constructs *LRX1:LRX1* and *LRX1:LRX1^ΔNT^*. Unlike the full-length *LRX1* which complements the *lrx1* mutant (Baumberger et al, 2001, Ringli 2010) and the *lrx1 lrx2* mutant (Suppl. Figure S2), *LRX1^ΔNT^* failed to induce wild-type root hairs in either of the mutants (Suppl. Figure S2).

### LRX1, LRX4, and LRX5 are high-affinity binding sites for RALF1

LRX4 has been shown to bind rapid alkalinization factor 1 (RALF1) a peptide hormone that also interacts with FER [15,42]. Here, we tested binding of RALF1 by LRX1. Transient expression of *LRX1*^ΔE^*-HA* and *RALF1-FLAG* in *N. benthamiana* followed by Co-IP and western blotting showed interaction of the two proteins (Figure 5A). This was confirmed by Y2H, where under selective conditions, yeast cells grew effectively in the presence of the two proteins (Suppl. Figure S3).

**Figure 5.**
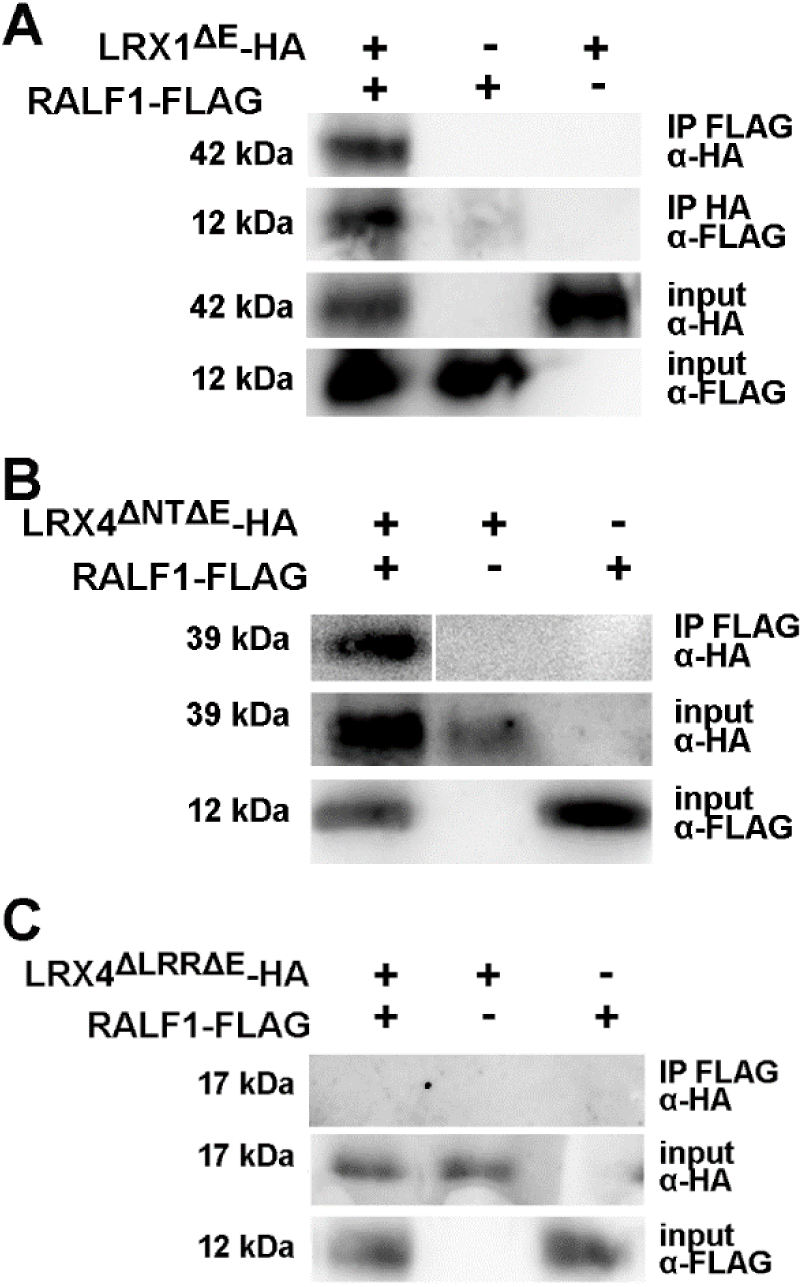
RALF1 is bound via the LRR domain of LRX proteins. (A) When co-expressed in tobacco LRX1^ΔE^ and RALF1 can be co-immunoprecipitated, indicative for interaction of the two proteins. (B and C) LRX4^ΔNTΔE^ (B) but not LRX4^ΔLRRΔE^ (C) are co-purified with RALF1, indicating that RALF1 is bound by the LRR domain. Antibodies used for IP and subsequent detection by western blotting are indicated.

The kinetics of the interaction of LRX proteins with RALF1 were tested with Biolayer Interferometry (BLITZ). The LRX^ΔE^-FLAG proteins of LRX1, LRX3, LRX4, and LRX5 used for this experiment were expressed transiently in tobacco. Expression of all proteins to comparable levels was confirmed by western blotting prior to BLITZ analysis. For RALF1, *in vitro* synthesized peptide was used. This analysis revealed a dissociation constant Kd of around 5 nM for the interaction of LRX1, LRX4, and LRX5 with RALF1 (Suppl. Figure S4). LRX3, by contrast, did not show interaction with RALF1 (Table 1).

**Table 1.**
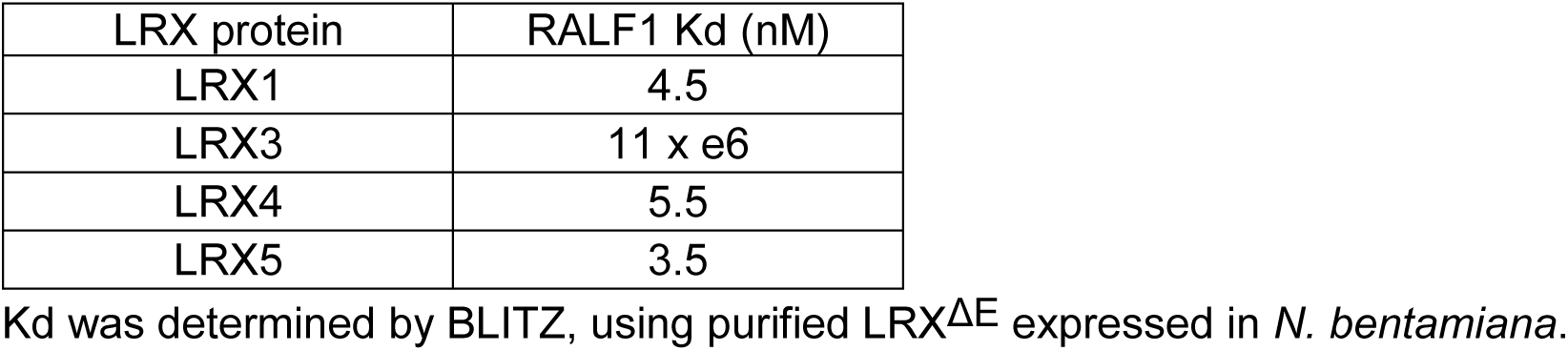
Dissociation constant of LRXs-RALF1 interactions

| LRX protein | RALF1 K<sub>d</sub> (nM) |
| --- | --- |
| LRX1 | 4.5 |
| LRX3 | 11 x e6 |
| LRX4 | 5.5 |
| LRX5 | 3.5 |

K<sub>d</sub> was determined by BLITZ, using purified LRX<sup>ΔE</sup> expressed in *N. benthamiana*.

Finally, the structural requirements for the LRX-RALF1 interaction was tested by expressing the *LRX4* deletion constructs *LRX4^ΔNTΔE^-HA* and *LRX4^ΔLRRΔE^-HA* with *RALF1-FLAG* in *N. benthamiana* for Co-IP experiments. As shown in Figure 5B and 5C, *LRX4^ΔNTΔE^-HA* but not *LRX4^ΔLRRΔE^-HA* showed co-purification with RALF1-FLAG, indicating that the LRR domain is necessary and sufficient for LRX-RALF1 interaction.

### Functional equivalence among different LRX proteins

Analyzes performed so far suggest that most LRX proteins have comparable functions while LRX3 appears to have overlapping but non-identical binding abilities [[38], this work]. To compare *in planta* the function and activity of *LRX* genes expressed in different tissues, trans-complementation experiments were performed. To this end, the genomic coding sequence of *LRX1* encoding a cmyc-tag at the beginning of the LRR domain that does not interfere with protein function [32] was cloned into an overexpression cassette containing the *35S* promoter and the resulting *35S:LRX1* construct was used for transformation of the *lrx345* triple mutant. Several independent homozygous transgenic lines were identified and characterized. Semi-quantitative RT-PCR confirmed expression of the transgene in the lines (Suppl. Figure S5A). For assessment of the complementation of the *lrx345* phenotype, alterations in plant growth and physiology were used as parameters. *lrx345* mutants grow smaller than the wild type both at seedling stage and at later stages when grown in soil [37]. This phenotype is alleviated in the transgenic lines (Figure 6A, Suppl. Figure S6A). The increased anthocyanin accumulation in *lrx345* mutant seedlings compared to the wild type is significantly reduced in transgenic lines (Figure 6B). The recently reported salt-hypersensitivity of the *lrx345* triple mutant resulting in reduced root growth and strong reduction in shoot growth in the presence of 100 mM NaCl [38] was also alleviated in the transgenic lines (Figure 6C and D). Hence, ectopically expressed *LRX1* can largely rescue the *lrx345* mutant phenotypes.

**Figure 6.**
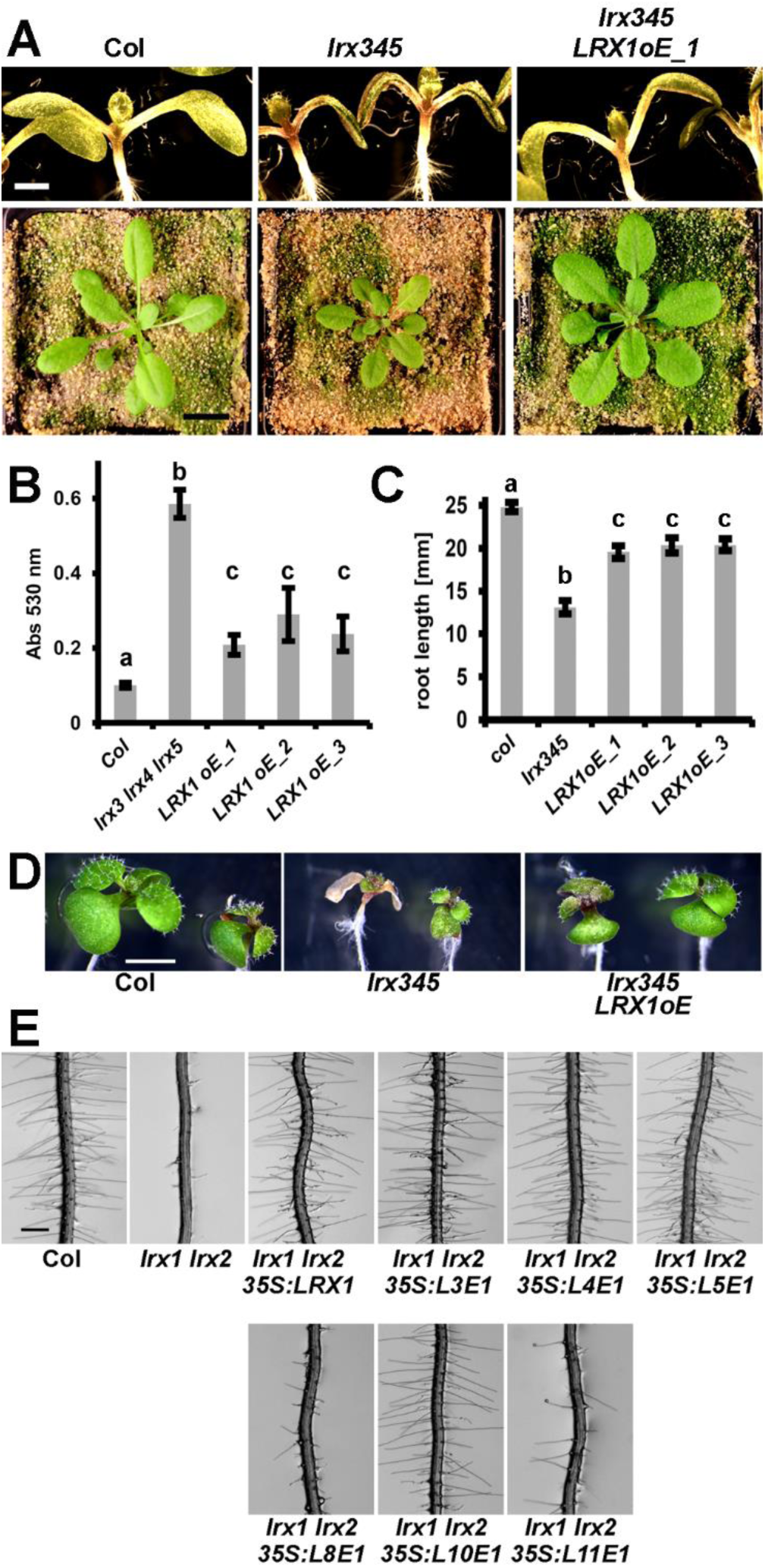
Functional redundancy among LRX proteins. (A-C) Complementation of the *lrx345* triple mutant with the *35S:LRX1* construct. Representative examples are shown. (A) Seedling shoots after 7 days of growth on half-strength MS (upper lane) and plants grown in soil (lower lane). (B) Anthocyanin accumulation in 12-days-old seedlings is increased in the *lrx345*. (C and D) The *lrx345* mutant seedlings grown in the presence of 100 mM NaCl show increased salt sensitivity, resulting in shorter roots (C) and impairment of shoot growth (D) compared to control Col, which are both alleviated by *LRX1* overexpression. Error bars represent SEM; different letters above the graphs indicate significant differences (T-test, N>20, P<0.05). (E) Complementation of the *lrx1 lrx2* root hair defect by chimeric construct of *L3,4,5,8,10*, and *11* (ATG start codon to CRD) fused to the extensin coding sequence of *LRX1* (*E1*) under the *35S* promoter. Representative examples of several independent transgenic lines for each construct are shown. Bars = 5 mm (A,D) and 0.5 mm (E).

In a complementary experiment, rescue of the intermediate root hair phenotype of the *lrx1* mutant, and the strong root hair phenotype of the *lrx1 lrx2* mutant [36] with *LRX3*, *LRX4*, and *LRX5* was tested. Due to the repetitive nature of the extensin coding sequences of *LRX3*, *LRX4*, and *LRX5*, these could not be stably maintained in *E.coli*. Therefore, and as previously described [37], the extensin-coding domains of *LRX3,4,5* genes were replaced by the one of *LRX1*. The resulting chimeric genes are referred to as *L3E1*, *L4E1*, and *L5E1* (L and E referring to the N-terminal moiety from the start codon to the CRD and the extensin coding sequence, respectively). The constructs encoding the chimeric proteins were placed under the control of the *35S* promoter and were transformed into the *lrx1* and *lrx1 lrx2* mutants. For each of the three constructs, several independent T2 lines were identified, all of which showed expression of the transgene (Suppl. Figure S5B). The root hair growth defect of the *lrx1* mutant (Suppl. Figure S6B) as well as the stronger *lrx1 lrx2* double mutant phenotype (Figure 6E) were suppressed by either of the three chimeric constructs.

The pollen-expressed *LRX8-LRX11* [34] form a separate phylogenetic clade and are more similar to pollen-expressed *LRXs* of other plants than to vegetative *LRXs* of Arabidopsis [29,48]. It was tested whether the N-terminal moieties of these LRXs are functionally comparable to LRX1. The chimeric constructs *L8E1*, *L10E1*, and *L11E1* under the *35S* promoter were transformed into the *lrx1 lrx2* double mutant. Several independent transgenic T2 lines were produced for each construct and transgene expression was confirmed in the lines (Suppl. Fig. S5C), but full rescue was only observed in plants expressing *L10E1*, while seedlings expressing *L8E1* or *L11E1* displayed poor root hair growth (Figure 6E). Expressing *L8E1* and *L11E1* in the *lrx1* single mutant resulted in rescue of the *lrx1* root hair phenotype (Suppl. Figure S6B), confirming that these proteins are in principle functional. These results suggest that the N-terminal moieties of different LRX proteins of vegetative tissues are sufficiently overlapping in their activities to replace *LRX* genes active in other vegetative cell types. By contrast, pollen-expressed *LRX*s have functionally diverged to varying degrees, some being similar but not equivalent to the *LRX*s expressed in vegetative tissues.

### LRX downstream signaling components

The FER homologs ANXUR1 and 2 (ANX1/2) are required to maintain pollen tube growth [6,49] and bind pollen-expressed RALF4 and 19 [21]. *AUN1^D94N^* and *MRI^R240C^* are two suppressors of the *anx1 anx2* male sterility [26,27]. To test whether the signaling pathways involving CrRLK1Ls and LRX proteins are comparable in reproductive and somatic tissues, the *lrx1 lrx2* double mutant was transformed with an *MRI:AUN1^D94N^-YFP* and a *MRI:MRI^R240C^-YFP* construct [26]. Several independent T2 plants transgenic for either of the two constructs displayed partial suppression of the *lrx1 lrx2* root hair phenotype (Figure 7, Suppl. Figure S7). Root hairs in *MRI^R240C^-YFP* transgenic lines are frequently shorter than in the wild type, which is attributable to the function of MRI^R240C^ [26]. This suggests that both AUN1 and MRI play a role in the signaling pathway downstream of LRX1 and LRX2.

**Figure 7.**
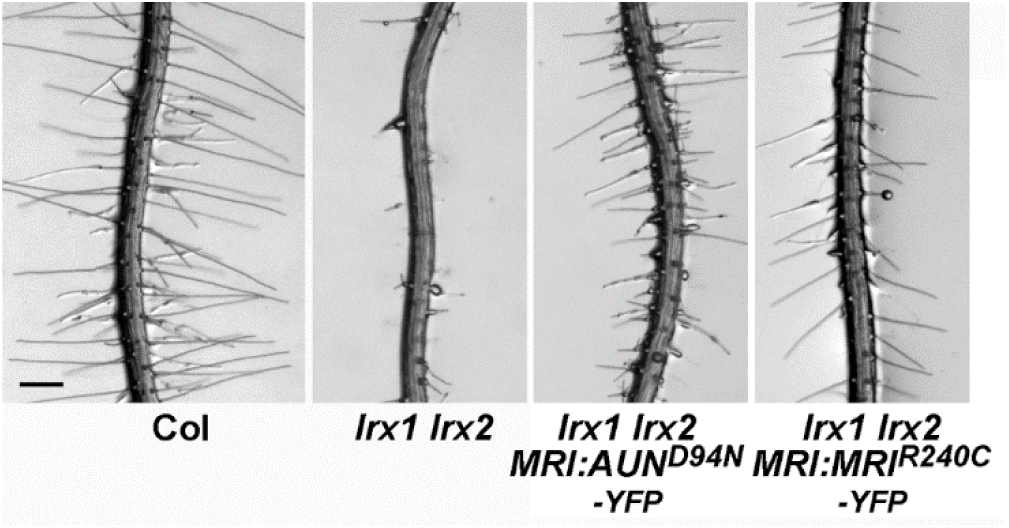
Expression of *AUN1^D94N^* and *MRI^R240C^* partially suppress the *lrx1 lrx2* root hair defect. Seedlings were grown for 5 days on half-strength MS in a vertical orientation. Representative examples of several independent transgenic *lrx1 lrx2* double mutant expressing either *pMRI:AUN1^D94N^-YFP* or *pMRI:MRI^R240C^-YFP* are shown. Bar = 0.5 mm

## Discussion

LRX proteins localize to the cell wall and have been shown to be involved in cell wall formation processes [34,36,37,39,40]. Only recently, their molecular function in this process has started to be unraveled by the identification of LRXs as extracellular binding sites of RALF peptides [35,38,41]. The association of LRX4 with the plasma membrane [34] and the identification of LRX4 as an interactor of the CrRLK1L-type receptor kinase FER revealed an LRX4-RALF1-FER signaling axis involved in regulating cell growth and vacuole development [35]. Support for this model is provided by the analysis of LRX4 deletion constructs, where a correlation between FER interaction and membrane association could be established. This signaling cascade thus also represents a link between the plasma membrane and a protein that is insolubilized in the cell wall via its extensin domain [31,32,33,50].

*FER* is expressed in many plant tissues and several *LRX* genes are expressed in different tissues with little overlap in their expression patterns [29]. Thus, it is plausible to assume that several LRXs of different tissues interact with FER to establish the FER-LRX interaction across diverse cell types. This assumption is corroborated by similar phenotypes of *fer-4* and an *lrx12345* quintuple mutant and by the demonstration of FER-RALF1-LRX1/2/4/5 interactions by different experimental approaches. *LRX6* and *LRX7* are two *LRX* genes not characterized so far. *LRX6* is expressed during lateral root formation and *LRX7* in flowers [29]. A heptuplicate *lrx* mutant line affected in all vegetatively expressed *LRX* genes would possibly show an even more severe defect in growth and development. Comparable to the interaction with FER, several LRXs including LRX1 interact with RALF1 and both interactions involve the LRR domain. This is corroborated by the very recently described LRX-RALF crystal structure [51]. Different LRXs bind an overlapping but not identical array of RALF peptides [35,38], and the full binding spectrum of the different LRXs might be even broader. One reason for the distinct RALF binding spectrum of LRXs is based on their expression pattern. Pollen-localized LRX8 shows a much higher affinity to the pollen-localized RALF4 than the root/shoot-localized RALF1 [41]. It can be speculated that the diverse affinities of RALFs and LRXs contribute to the specificities of the plant’s response to different RALF peptides that have distinct biological activities [17,18,19,20,25,52]. An additional regulatory layer is added by the pH that is influenced by e.g. RALFs and modifies their binding to interaction partners [20]. Crystallization analyses suggest that LRXs form dimers and both monomers can bind a RALF peptide [51]. Whether LRXs form homo- and/or heterodimers in vivo and whether these can bind different RALFs remains to be unraveled. Possibly, the exact pairs of LRX-RALF interactions influence binding of other proteins. These might include additional CrRLK1Ls but possibly also other plasma membrane-localized receptors. BAK1 is a co-receptor binding other receptor kinases such as FER or FLS2 [53] but also RALF1 and is required for the response of the plant to RALF1 [25]. It will be interesting to investigate whether some LRXs can interact with BAK1 or, alternatively, with apoplastic proteins not associated with the plasma membrane. These lines of research need to be followed to better understand the function of LRX proteins in cell wall development.

Earlier experiments with the root hair-expressed *LRX1* and *LRX2* suggested synergistic interaction and functional equivalence of these two LRXs [36]. The rescue experiments for *lrx1*, *lrx1 lrx2*, and *lrx345* mutants suggest that LRXs of vegetative tissues are functionally similar, as all combinations of mutants with these genes resulted in rescue of the mutant phenotypes. In contrast, the pollen-expressed *LRX8* and *LRX11* appear to have too strongly diverged to fulfill the same ‘vegetative’ functions as they barely rescue the *lrx1 lrx2* double mutant. This divergence is in part supported by phylogenetic analyses that revealed evolutionary separation of pollen- and vegetatively expressed LRXs [29,48]. The rescue experiments have also revealed that the construct *35S:L3E1* complements the *lrx1* and *lrx1 lrx2* mutants, which is remarkable since LRX3 fails to interact with FER [also reported by [38]] and RALF1. LRX proteins possibly interact with different, so far unknown proteins in addition to FER and the identified RALFs, and this activity might be sufficient for complementation of the *lrx* mutant phenotypes. Alternatively, the *in vivo* binding activity of LRX3 is not identical to the one observed in yeast-two-hybrid, BLITZ, or Co-IP experiments of tobacco-expressed proteins. Future experiments on binding capacities of different LRXs to so far unknown proteins or other CrRLK1Ls will be necessary to clarify this issue. In pollen tubes, the *FER*-homologs *ANX1*/*2* and *BUPS1/2* but not *FER* are expressed [6,10,21]. Thus, pollen-expressed LRXs possibly interact with these CrRLK1Ls. The finding that expression of *AUN1^D94N^*, a dominant hypomorph variant of AUN1 partially suppresses not only *anx1 anx2* but also the *lrx8-lrx11* quadruple mutant pollen bursting phenotype [27] is indicative of LRX and ANX1/ANX2 being active in the same pathway. The *anx1 anx2* suppressor mutants *AUN1^D94N^* and *MRI^R240C^* also partially suppress the *lrx1 lrx2* mutant root hair phenotype. Hence, the LRX-RALF-CrRLK1L signaling pathway is at least partially conserved among different cell types, suggesting that CrRLK1Ls use a common set of downstream signaling components.

Protein-protein interactions analyzed so far involve the LRR domain of LRXs. Yet, the NT-domain of unknown function is also important for LRX activity, since an NT deletion construct in LRX1 impairs protein function. The development of a dominant negative effect when expressing an extensin-less LRX1 [32,33] also depends on the presence of the NT-domain, even though binding of RALF1 and FER are NT-domain independent. Again, these data indicate possible additional features of LRX proteins beyond their interaction with CrRLK1Ls and RALFs that contribute to their full biological activity. The activity of the NT-domain and whether it is involved in binding other proteins remains to the determined.

Potential differences in the function of the LRX extensin domains have not been investigated here, but rather avoided by using the *LRX1* extensin coding sequence for all complementation constructs. The extensin domain is variable among the LRXs both in terms of length and the repetitive motifs typical for this structural protein domain [29,31,50]. It is possible that the extensin domains of the different LRXs have adapted to the specific cell wall composition of the various tissues they are active in, which would explain the considerable differences in the extensin domains [29]. Extensins form covalent links with other extensins or with polysaccharides in the cell wall [50,54,55], and the composition of cell walls differs considerably among cell types [56,57].

Cell growth requires the controlled simultaneous expansion of the cell wall and the protoplast, and this might be monitored through a FER-LRX interaction that depends on physical proximity of the two proteins and, consequently, of the plasma membrane and the cell wall. The observed regulation of vacuolar dynamics required for cell growth by FER and LRXs [42], supports this hypothesis, since both partners are attached/embedded in their subcellular structure. Detaching LRX proteins from the cell wall by removal of the extensin domain interferes with this balanced system, causing a defect in cell growth [32,33,35]. This work expands the FER-LRXs interaction to LRX5 in root/shoot tissue and reveals an LRX1-RALF1-FER interaction network important for proper root hair growth. The functional redundancy among LRX proteins of different vegetative and reproductive tissues indicates that LRXs function is not limited to interaction with FER. Clearly, different RALFs and probably different *Cr*RLK1Ls are demonstrated or potential binding partners of LRXs, where the specificities of interaction might reflect differences in the biological processes triggered by the interactions. In future studies, it will be important to analyze the dynamics of LRX-RALF-FER interactions and to identify additional intra- and extracellular factors involved in the process to better understand the implications and mechanisms of this cell wall integrity sensing network in the regulation of cell growth.

## Materials and Methods

### Plant growth and propagation

*Arabidopsis thaliana* of the ecotype Columbia were used for all experiments. Seeds were surface sterilized with 1% Sodium hypochlorite, 0.03% TritonX-100, washed three times with sterile water, and, unless stated otherwise, plated on half-strength MS plates (0.5X MS salt, 2% Sucrose, 0.5 mg/L nicotinic acid, 0.5 mg/L pyridoxine-HCl, 0.1 mg/L thiamine-HCl, glycine 2 mg/L, 0.5 g/L MES, pH 5.7, 0.6% Gelzan (Sigma); referred to as standard medium) under a 16hrs light – 8 hrs dark photoperiod at 22°C. For propagation and crossings, plants were grown under the same conditions in soil.

For selection of transgenic lines, seeds of plants used for Agrobacterium (GV3101)-mediated transformation by the floral-dip method, were plated on 0.5 MS plates, 2% sucrose, 0.8% bactoagar, supplemented with appropriate antibiotics; 100 μg/ml ticarcillin, 50 μg/ml kanamycin or 10 μg/ml glufosinate-ammonium.

### Molecular markers

PCR-based molecular markers used to produce the different lines are described in [37] for *lrx3*, *lrx4*, and *lrx5*, [58] for *lrx1*, [36] for *lrx2*, and [42] for *fer-4*. The primers used for the PCR reactions are listed in the Supplementary Data Table S1.

### DNA constructs

For the *LRX4^ΔE^* construct, the coding sequence of the N-terminal half of LRX4 was amplified using the primers LRX4oE_XhoI_F and LRX4_PstI_R (Supplementary Table S2). This product was digested with *Xho*I and *Pst*I and ligated with a fragment encoding a double FLAG tag with a *Pst*I and a *Xba*I site at the 5’ and 3’ end, respectively, into the plasmid pART7 [59] digested with *Xho*I and *Xba*I, resulting in the *35S:LRX4^ΔE^-2FLAG* construct. All final constructs were control sequenced.

For *LRX4^ΔLRRΔE^*, the sequence from the start codon to the end of the NT-domain was amplified with LRX4oE_XhoI_F and LRX4*^ΔLRR^*_PstI_R, the resulting fragment digested with *Xho*I and *Pst*I and cloned into the plasmid *35S:LRX4^ΔE^-2FLAG* cut with the same enzymes. For *LRX4^ΔNTΔE^*, the sequence encoding the signal peptide was amplified with primers LRX4_XhoI_F and LRX4_ΔNT_R and the LRR domain with the primers LRX4_ΔNT_F and LRX4_PstI_R, the fragments were digested with *Xho*I/*Bam*HI and *Bam*HI/*Pst*I, respectively and cloned by triple ligation into *35S:LRX4^ΔE^-2FLAG* cut with *Xho*I and *Pst*I.

For *LRX1^ΔE^ -FLAG*, the *LRX1* fragment was amplified using the primers LRX1_XhoI_F and LRX1_PstI_R, digested with *Xho*I/*Pst*I and cloned into the vector *pART7_LRX4^ΔE^-FLAG* digested with the same enzymes to release the *LRX4^ΔE^* sequence. For the *35S:L1E1* construct, the plasmid *35S:LRX1^ΔE^-2FLAG* was opened with *Pst*I and *Xba*I and a *Pst*I-*Spe*I fragment containing the extensin-coding sequence [36] was inserted.

The *LRX1:LRX1^ΔE^* construct containing the *cmyc* tag in front of the *LRR* domain is described elsewhere [33]. For the *LRX1:LRX1^ΔLRRΔE^* construct, the promoter and coding sequence up to the end of the cmyc-tag was amplified with the primers LRX1_Prom1000_F and LRX1_ΔLRR_SpeI_R, and the resulting fragment was digested with *Mlu*I (in the *LRX1* promoter) and *Spe*I (at the end of the myc tag sequence) and cloned into the *LRX1:LRX1* construct cut with the same enzymes (*Spe*I overlapping with the stop codon of the *LRX1* coding sequence). For *LRX1:LRX1^ΔNTΔE^*, the promoter and signal peptide coding sequence was amplified with the primers LRX1_Prom1000_F and LRX1_ΔNT_SaII_R and the resulting fragment was digested with *Mlu*I (in the promoter) and *Sal*I (at the end of the signal peptide sequence) and cloned into *LRX1:LRX1^ΔE^* cut with the same enzymes (the *Sal*I site in the *LRX1:LRX1^ΔE^* construct is at the beginning of the *cmyc* coding sequence). For *LRX1:LRX1^ΔNT^* the *Mlu*I-*Sal*I fragment of *LRX1:LRX1^ΔNTΔE^* was ligated into *LRX1:LRX1* cut with the same enzymes.

For the *35S:L3/4/5/8/10/11-E1* constructs, the coding sequences from the ATG to the CRD-coding sequence were amplified with primers (Suppl. Table S2) introducing a *Kpn*I or an *Xho*I and a *Pst*I site at the 5’ and 3’ end of the PCR product, respectively, and the fragments were ligated into *35S:L1E1* cut with the same enzymes to release the L1 coding sequence.

All the *pART7*-based expression cassettes were cut out with *Not*I and cloned into the binary vector *pART27* [59] cut with the same enzyme.

Cloning of *MRI^R240C^* CDS without stop codon in Gateway® compatible binary vector *pMRI:-:GW-YFP* plasmid (*pABD83*, Basta Resistance) to obtain *pMRI:AUN1^D94N^-YFP* was described previously (Boisson-Dernier et al., 2015). To obtain *pMRI:AUN1^D94N^-YFP, AUN1^D94N^* CDS without stop codon in *pDONR207* (Invitrogen) (Franck et al., 2018b) was remobilized into *pABD83*.

The *BD-LRX4* and *AD-FER^ECD^* constructs for the Yeast-two-hybrid experiment were cloned as previously described [35], where *NtermFER* equals *AD-FER^ECD^* and *LRR4* equals *BD-LRX4*. For the *BD-LRX1/2/3/5* constructs, the coding sequence of the LRR domain coding sequence of the *LRXs* was amplified using primers (Suppl. Table S2) to introduce a *Bam*HI and a *Xho*I site at the 5’ and 3’ end of the PCR fragments, respectively. These were cloned into *pJET1.2* (Thermo Scientific) and correct clones were cut with *Bam*HI and *Xho*I and ligated into *pGBKT7* cut with *Bam*HI and *Sal*I. The AD-RALF1 construct was cloned into *pJET1.2* (Thermo Scientific) by amplification of the coding sequence with the primers y2h_RALF1_F and y2h_RALF1_R. A correct clone was cut with *Eco*RI and *Xma*I and ligated into *pGADT7* cut with *Eco*RI and *Xma*I.

### Phenotyping of seedling growth properties

For the quantification of gravitropism, seedlings were grown in a vertical orientation on standard MS medium for 8 days, and the ratio of root progression in the vertical axes over total root length was used as the parameter, as described [47]. For measurements, the plates were scanned and analyzed by ImageJ. To ascertain consistent results, seedlings of different generations were used and at least 10 seedlings were measured for one data point.

The accumulation of anthocyanin was quantified on 12 days-old seedlings grown in a vertical orientation on standard medium by published methods [60,61]. Twenty seedlings were pooled and incubated in 45% Methanol, 5% acetic acid. After centrifugation for 5 min at RT and 13’000 rpm, the supernatant was used to measure absorption at 530 nm for anthocyanin and at 657 nm for chlorophyll content correction; final value =Abs_530nm_-(0.25xAbs_657nm_). One data point in the graph is the average of quadruplicates.

For root length measurements, seedlings were grown for 7 days on standard medium in a vertical orientation, plates were scanned, and ImageJ was used to measure root length. The average of at least 15 seedlings was used for one data point.

Root hair phenotypes were assessed in 5 days-old seedlings grown in a vertical orientation on standard medium. Pictures of root hair were taken with a MZ125 Binocular (Leica), using a DFC420 digital camera (Leica).

### Co-immunoprecipitation

Co-IP experiments were performed exactly as previously described [35]. For pulldown and co-IP analysis of the different constructs indicated in the experiments were infiltrated into *Nicotiana benthamiana* leaves, and after 48 hrs, the leaves were excised and grinded in liquid nitrogen. The tissue powder was re-suspended in extraction buffer [200 mM Tris-HCl (pH 7.5), 150 mM NaCl, 1 mM DTT, 1 mM PMSF, protease inhibitor and 0.5% Triton X-100]. The suspension was incubated on ice for 20 minutes and then centrifuged at 13,000 rpm for 30 minutes at 4°C. The supernatant obtained was then incubated with GFP-trap agarose beads, anti-HA, or anti-FLAG magnetic beads overnight at 4°C on a rotating shaker. After incubation, the beads were washed three times with the wash buffer (extraction buffer containing 0.05% Triton X-100) and boiled in SDS-PAGE loading buffer for 15 minutes at 75°C. The immunoprecipitates were then run on a 10% SDS-PAGE and transferred to nitrocellulose membrane to perform Western blotting.

### BLITZ analysis

The BLITZ experiments were performed as previously described [41].

The *LRR^ΔE^*–FLAG versions of the different *LRXs* were expressed under the *35S* promoter in *N. benthamiana*, presence of proteins was checked by western blotting, and proteins were immune-precipitated as described above. After immunoprecipitation, elution was performed with 30 μl of 1M Glycine (pH 2.0) buffer for 2 min in a Thermomixer (Eppendorf) at 1200 rpm, then beads were spun down for 2 min at 1300xg at RT, and the supernatant was neutralized with 30 μl of 1M Tris-HCl (pH 9.5). Protein concentration was determined by Qubit measurement (Quant-iTTM Protein Assay kit, Invitrogen). Samples were diluted 1:1 with sample diluent buffer (Pall FortéBio cat18-1091) to a concentration of 0.142 mg/ml for analysis using the BLITzⓇ system. The same buffer was used to dilute the anti-FLAG M2 antibody (Sigma-Aldrich) 1:50 to a final concentration of 4 μg/ml. A 1:1 mix of sample:antibody was then incubated for 30 minutes at RT, and loaded onto the protein A biosensor (Pall FortéBio cat 18-5010). The experiment was divided into 5 different steps: Initial baseline duration (30 s), Loading duration (120 s), Baseline duration (30 s), Association duration (120 s), and Dissociation duration (120 s). Different RALF synthetic peptide concentrations (200 μM, 150 μM, 100 μM, 50 μM, 20 μM, 15 μM, 10 μM, 4 μM, 2 μM and 0.2 μM) were added to quantify the protein interaction.

### Western blotting

To test the accumulation of LRX1^ΔE^, LRX1^ΔLRRΔE^, and LRX1^ΔLNTΔE^ proteins, root material of 300 seedlings grown for 10 days in a vertical orientation was collected and ground in liquid N2. Around 50 mg of fresh material was extracted with 200 μl 0.1% SDS by vortexing, immediately followed by heating to 95°C for 5 min. After cooling, material was centrifuged at 13’000 rpm for 10 min and 20 μl of the supernatant was used for SDS-PAGE and blotting to nitrocellulose membranes using semi-dry blotting. After over-night blocking of the membranes in 1xTBS, 0.1% Tween-20, 5% low-fat milk powder, the membranes were incubated in 1xTBS, 0.1% Tween-20, 0.5% low-fat milk powder containing primary antibodies as indicated in the figures, followed by a peroxidase-coupled secondary antibody, diluted 1:1000 each. After each antibody incubation, the membranes were washed three times with the antibody-incubation solution. The signal of the secondary antibody was detected using the ECL technology.

### RT-PCR

Semi-quantitative RT-PCR was performed on RNA isolated of 10 days-old seedlings using the total RNA isolation system (Promega). Reverse transcription was performed on 300 ng of total RNA using the iScript advanced kit (BioRad). PCR was performed using gene-specific primers as listed in the supplementary data Table S3. Correct amplification of the expected DNA band was verified by sequencing of the PCR products.

### Yeast-two-hybrid

Transformation of the yeast strain PJ69-4A [62] was done following standard procedures and quadruple drop-out medium lacking Leu, Trp, His, and Ade were used to screen for positive interactions after 4 days incubation at 30°C. Always three different colonies containing both vectors were mixed and plated in triplicates on quadruple drop-out medium.

### Membrane fractionation

Membrane fractionation was performed as described [34] using an established method [63]. Homogenized tissue samples were suspended in 3 volumes of ice-cold extraction buffer [250 mM sorbitol; 50 mM Tris-HCl, 2 mm EDTA; pH 8.0 (HCl); immediately before use add: 5 mM DTT; 0.6 % insoluble PVP; 0.001 M PMSF; 10 µL/mL Protease Inhibitor Cocktail (Sigma P9599)]. The material was first centrifuged at 5,000g and 10,000g for 5 minutes each at 4°C to remove cell debris. The supernatant was then centrifuged at 40,000 rpm for 1 hour at 4°C and the pelleted membrane fraction was resuspended in [5 mM KH2PO4; 330 mM sucrose; 3 mM KCl; pH 7.8 (KOH); 0.5% n-Dodecyl-β-D-maltopyranoside]. The samples were used for SDS-PAGE and Western blotting, where the LRX1^ΔE^ LRX1^ΔLRRΔE^ were detected with an anti-cmyc and LHC1a with an anti-LHC1 antibody.

### Gene identifiers of genes used in this study

FER: At3G51550; RALF1:At1G02900; LRX1: At1g12040; LR2: At1g62440; LRX3: At4g13340; LRX4: At3g24480; LRX5: At4g18670; LRX8: At3g19020; LRX10: At2g15880; LRX11: At4g33970

## Acknowledgments

We are grateful to Ueli Grossniklaus and Valeria Gagliardini for introducing SG to the BLITZ experimental procedure. This work was supported by the Swiss National Science Foundation (grant Nr. 31003A_166577) to CR and in part by a grant from the University of Cologne Centre of Excellence in Plant Sciences to ABD.

## Supplementary Data

Suppl. Figure S1 Interaction of LRXs and FER.

Suppl. Figure S2 LRX1^ΔNT^ does not complement the *lrx1* and *lrx1 lrx2* mutants.

Suppl. Figure S3 Interaction of LRX1 and RALF1 in yeast.

Suppl. Figure S4 BLITZ output data.

Suppl. Figure S5 RT-PCR confirming expression of the transgenes.

Suppl. Figure S6 Complementation of *lrx345* and *lrx1* mutants.

Suppl. Figure S7 Fluorescence of *AUN1^D94N^-YFP and MRI^R240C^-YFP* transgenic lines.

Suppl. Table S1 Primers used for genotyping.

Suppl. Table S2 Primers used for cloning.

Suppl. Table S3 Primers used for RT-PCR.

