## Supplementary Figures for "LRR-extensins of vegetative tissues are a functionally conserved family of RALF1 receptors interacting with the receptor kinase FERONIA"

### Suppl Figure S1

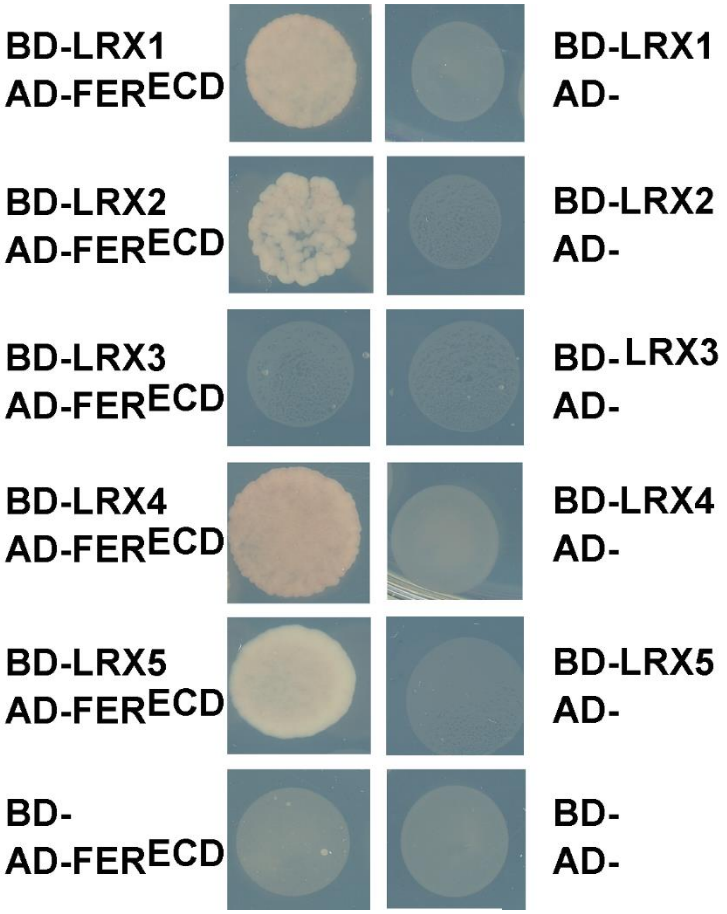


**Suppl. Figure S1** Interaction of LRXs and FER.

The yeast strain PJ69-4A was transformed with *pGADT7-FER*$^{ECD}$ and *pGBKT7-LRRs* (encoding the LRR domain of the different LRXs as indicated), and with either of the constructs with empty plasmids for the negative controls. The clear difference in yeast growth under selective conditions (-Trp, Leu, His, Ade drop-out medium containing 5 mM 3-AT) indicates interaction of the LRX proteins with FER.

### Suppl Figure S2

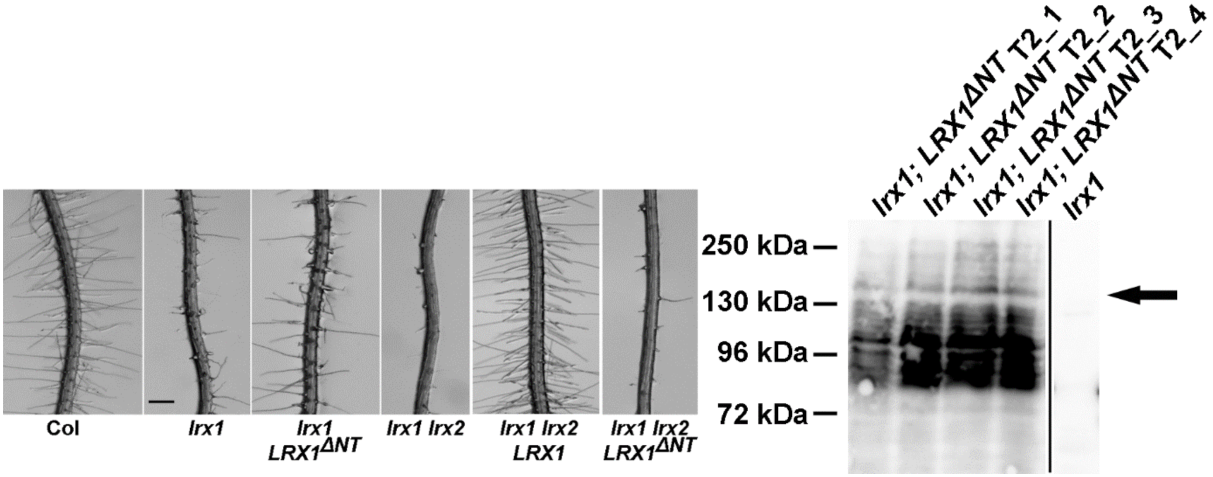


**Suppl. Figure S2** LRX1$^{\Delta NT}$ does not complement *lrx1* and *lrx1 lrx2* mutants.

Expressing *LRX1$^{\Delta NT}$* under the *LRX1* promotor fails to produce wild type-like root hair development in the *lrx1* or *lrx1 lrx2* double mutant. The full-length LRX1 induces wild-type root hairs in the *lrx1 lrx2* double mutant. Western blot demonstrating expression of the recombinant protein in the transgenic lines. Arrow indicates expected band of LRX1$^{\Delta NT}$. Bar: 0.5 mm

### Suppl Figure S3

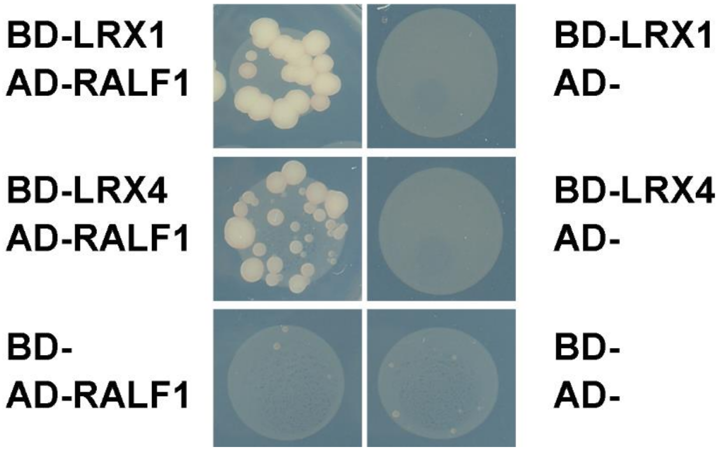


**Suppl. Figure S3** Interaction of LRX1 and RALF1 in yeast.

The yeast strain PJ69-4A was transformed with *pGADT7-RALF1* and *pGBKT7-LRR1* or *pGBKT7-LRR4*, and with either of the constructs with empty plasmids for the negative controls. The clear difference in yeast growth under selective conditions (Trp, Leu, His, Ade drop-out medium) indicates interaction of the LRX proteins with RALF1.

### Suppl Figure S4

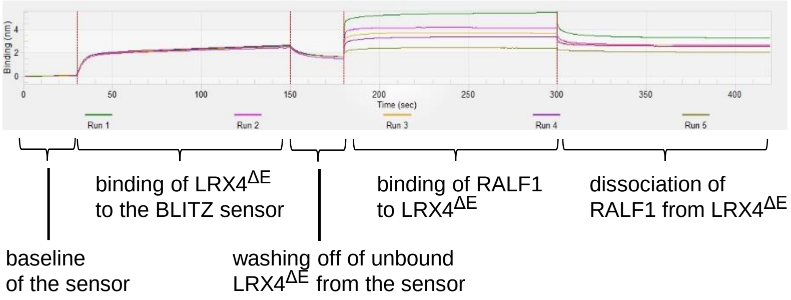


**Suppl. Figure S4** BLITZ output data.

Tobacco expressed and immunoprecipitated LRX4$^{\Delta E}$ –FLAG is attached to the sensor resulting in a signal for bound protein. After washing to remove excess LRX4$^{\Delta E}$ –FLAG protein. RALF1 peptide is added in different concentrations, followed by dissociation of RALF1. Association/dissociation or RALF1 depends on the concentration of applied RALF1, based on which the Kd is calculated.

### Suppl Figure S5

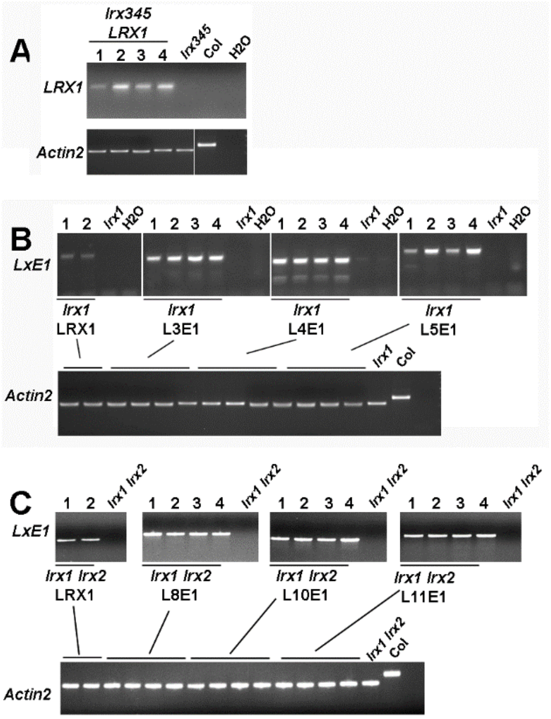


**Suppl. Figure S5** RT-PCR confirming expression of the transgenes.

Total RNA was extraceted from 7-days old seedlings and transgene-specific RT-PCR was performed to confirm expression of the transgenes. In all experiments, *Actin2* was used as endogenous control for comparable amounts of RNA in all samples and absence of contaminating genomic DNA. Genomic DNA of *Actin2* results in a longer PCR product as shown.

(A) Expression of *LRX1cmyc* in the *lrx345* triple mutant background.

(B) and (C) the label *LxE1* referring to the different transgenes as indicated. the lane «Col» represents genomic DNA.

### Suppl Figure S6

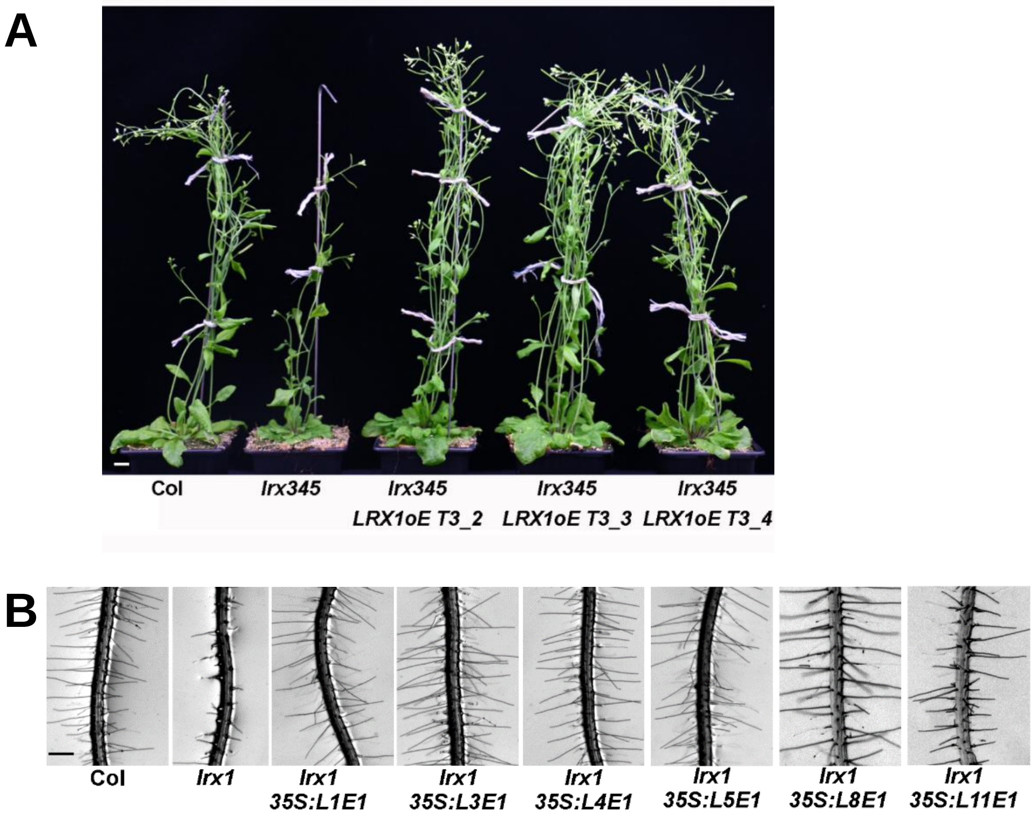


**Suppl. Figure S6** Complementation of the *lrx345* and *lrx1* mutants.

(A) *lrx345* triple mutant plants are smaller at flowering stage than the wild type (Col). This phenotype is complemented by the *35S:LRX1cmyc* construct.
(B) The *lrx1* root hair defect is complemented by the *LRX* chimeric constructs.
Bar: 1cm (A); 0.5 mm (B,C)

### Suppl Figure S7

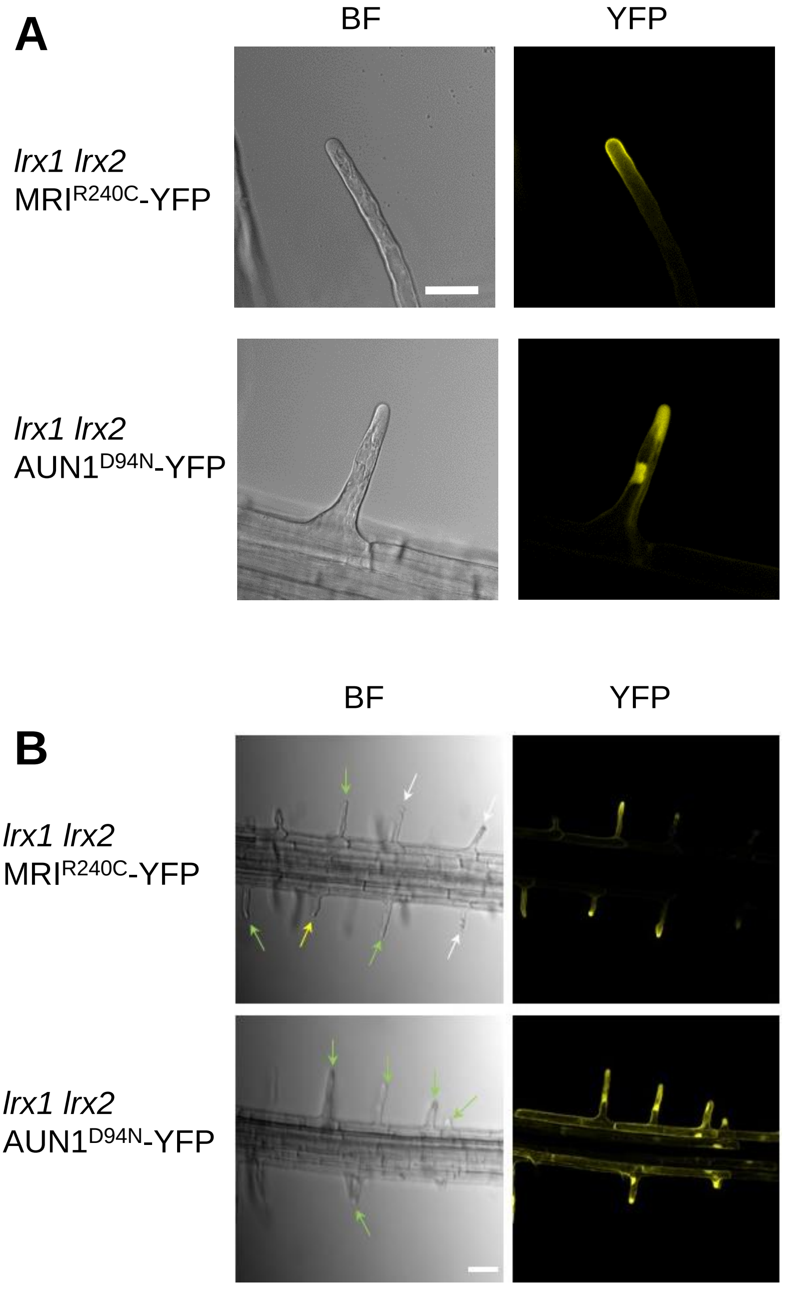


**Suppl. Figure S7** Fluorescence *of AUN1$^{D94N}$* -YFP and *MRI$^{R240C}$-YFP* transgenic lines.
Transgenic lines expressing *MRI$^{R240C}$-YFP* or *AUN1$^{D94N}$ –YFP* show YFP fluorescence in the cytoplasm and the plasma membrane, respectively. Individual root hairs (A) and entire roots (B) are shown. (B) When expressing *MRI$^{R240C}$-YFP* root hair growth is frequently arrested resulting in shorter root hairs (yellow arrows) whereas others grow normal (green arrows). Some root hairs still burst after some time (white arrow). Bar=20 µm (A), 50 µm (B).
