## Supplementary Tables for "LRR-extensins of vegetative tissues are a functionally conserved family of RALF1 receptors interacting with the receptor kinase FERONIA"

Supplementary Table S1 Primers used for genotyping

| Target Sequence | Primer | Sequence | |
|---|---|---|---|
| ***lrx1*** | LRX1_2800_F | GAAATCGATCCCGAGTCGTTG | *Bsi*EI digests the *lrx1* mutant |
| | LRX1_2980_R | TGGAGAGAAGATGTTGTAACATC | |
| ***lrx2*** | lrx2-1_F | TTGCGGATAATGATCTTGGAGGTTG | *Bcc*I digests the wild type |
| | lrx2-1_R | CTGCTTGGTATAACTCCTGTGAATC | |
| ***LRX3*** | LRX3_seq_F | TACTGTACCACACCGGTTTAAC | |
| | LRX3_LRR_R | AATTCTAGATCAACGGCCACAACTAAACGATC | |
| ***lrx3*** | LRX3_seq_F | TACTGTACCACACCGGTTTAAC | |
| | LBb1 | GCGTGGACCGCTTGCTGCAACT | |
| ***LRX4*** | LRX4_seq_F | CCATAACCGGTTCCGGTTTGAG | |
| | LRX4_3'b | TTTAACAACAGAACGACCACAAC | |
| ***lrx4*** | LRX4_seq_F | CCATAACCGGTTCCGGTTTGAG | |
| | Gabi_Lb | GGGAATGGCGAAATCAAGGCATCG | |
| ***LRX5*** | LRX5_F2 | GCTTGGTTTGTTAACAGATCTC | |
| | LRX5_3' | GCCGGAATCTTACCAGAGAATC | |
| ***lrx5*** | LRX5_F2 | GCTTGGTTTGTTAACAGATCTC | |
| | LBb1 | GCGTGGACCGCTTGCTGCAACT | |
| ***FER*** | FER_F1 | GATTACTCTCCAACAGAGAAAATCCT | |
| | FER_R1 | CGTATTGCTTTTCGATTTCCTA | |
| ***fer-4*** | FER_F2 | ACGGTCTCAACGCTACCAAC | |
| | FER T-DNA | TTTCCCGCCTTCGGTTTA | |

Supplementary Table S2 Primers used for cloning

| Primer | Sequence |
|---|---|
| LRX1_XhoI_F | CTCGAGTTAGTAAAATTGGGTTTCTTGAC |
| LRX1_PstI_R | CTGCAGTCCACAGGGCGAGCAAG |
| LRX1_ΔLRR_SpeI_R | ACTAGTTCAGGGTACCGAATTCAAGTCCTC |
| LRX1_ΔNT_SalI_F | GTCGACCGAAAACCCGAGTCGTTGCTGGC |
| LRX1_ΔNT_SalI_R | CCGTCGACCATCGTCGTGGTCTGCTTTG |
| LRX1_Prom1000_F | AAAGTGAGGTATTTAGGTCATT |
| LRX4oE_XhoI_F | GGTACCGTATCGTATCGTGAAATGAAGAAC |
| LRX4_PstI_R | CTGCAGTCCACCGAAGGCCGTG |
| LRX4_ΔLRR_PstI_R | GTTCTGCAGTTCCGGTTATCAAGAGCTTTAG |
| LRX4_ΔNT_F | CTAGGATCCAAAGCTCTTGATAACCGGAAG |
| LRX4_ΔNT_R | CTAGGATCCTGAGATTGAGAGAGAATGAGAG |
| Yeast two hybrid: | |
| LRX1_BamHI_F | GGATCCCTTCTCCTTCATACCCGAAAAC |
| LRX1_XhoI_R | CTCGAGTCAACTGCAATCCACAGGGCGAG |
| LRX2_BamHI_F | CAGGGATCCCTTCTCCTTCTAGTCCGAAAAC |
| LRX2_XhoI_R | GTCCTCGAGCTAACTACAATCAACAGAGCGTGAG |
| LRX3_BamHI_F | GGATCCTGGCGCTTGATAATCGGAAG |
| LRX3_ XhoI_R | ACGCTCGAGTCATCCACAATTTACCGGCGGAC |
| LRX5_BamHI_F | AGCGGATCCCGGCTTTAGATAACCGCCGAATCC |
| LRX5_XhoI_R | AGTCTCGAGTCAAAAGGAACCACAATCCACCG |
| RALF1_EcoRI_F | GAATTCAATAGAAGAATATTGGCGAC |
| RALF1_SmaI_R | CCCGGGTCAACTCCTGCAACGAATTTTGC |
| LxE1 constructs: | |
| LRX3_XhoI_F | CTCGAGATGAAGAAGACGATTCAAATC |
| LRX3_PstI_R | CTGCAGTTTACCGGCGGACGAGAC |
| LRX4_XhoI_F | GGTACCGTATCGTATCGTGAAATGAAGAAC |
| LRX4_PstI_R | CTGCAGTCCACCGAAGGCCGTG |
| LRX5_XhoI_F | CTCGAGATGAAGACGAAGATGATGATG |
| LRX5_PstI_R | TGACTGCAGCCACAATCCACCGGCGGAAGAG |
| LRX8_XhoI_F | TGACTCGAGATGACCCGAAGAACAATGGAG |
| LRX8_PstI_R | TGACTGCAGCTGCAATCCACAGGACGGC |
| LRX10_KpnI_F | AGAGGTACCGTTCCAAGTCATGACTAAACCTCC |
| LRX10_PstI_R | GATCTGCAGCTACAATCAACTGGACGGCTAATC |
| LRX11_XhoI_F | AGACTCGAGATGCCCTTCTATAAGCAGCC |
| LRX11_PstI_R | TCTCTGCAGCTACAATCAACCGGGCGGTTGCTA |

Supplementary Table S3 Primers used for RT-PCR

| Target Sequence | Primer | Sequence |
|---|---|---|
| ***L1E1*** | LRX1 cmyc F | GAAATCGATCCCGAGTCGTTG |
| | LRX1_Ext_R | TGGAGAGAAGATGTTGTAACATC |
| ***L3E1*** | LRX3_seq_F | TACTGTACCACACCGGTTTAAC |
| | LRX1_Ext_R | TGGAGAGAAGATGTTGTAACATC |
| ***L4E1*** | LRX4_seq_F | CCATAACCGGTTCCGGTTTGAG |
| | LRX1_Ext_R | TGGAGAGAAGATGTTGTAACATC |
| ***L5E1*** | LRX5_F2 | GCTTGGTTTGTTAACAGATCTC |
| | LRX1_Ext_R | TGGAGAGAAGATGTTGTAACATC |
